# Nanopore sequencing enables comprehensive transposable element epigenomic profiling

**DOI:** 10.1101/2020.05.24.113068

**Authors:** Adam D. Ewing, Nathan Smits, Francisco J. Sanchez-Luque, Jamila Faivre, Paul M. Brennan, Seth W. Cheetham, Geoffrey J. Faulkner

## Abstract

We apply long-read nanopore sequencing and a new tool, TLDR (Transposons from Long Dirty Reads), to directly infer CpG methylation of new and extant human transposable element (TE) insertions in hippocampus, heart, and liver, as well as paired tumour and non-tumour liver. Whole genome TLDR analysis greatly facilitates studies of TE biology as complete insertion sequences and their epigenetic modifications are readily obtainable.

## Main

Transposable elements (TEs) pervade our genomic architecture and broadly influence human biology and disease^1^. Recently, Oxford Nanopore Technologies (ONT) long-read DNA sequencing has enabled telomere-to-telomere chromosome assembly at base pair resolution, including of high copy number TEs previously refractory to short-read mapping^2–4^. While most evolutionarily older TEs have accumulated sufficient nucleotide diversity to be uniquely identified, recent TE insertions are often indistinguishable from their source elements when assayed with short-read approaches.

Each diploid human genome contains 80-100 potentially mobile long interspersed element 1 (LINE-1) copies, referred to here as L1Hs (L1 Human-specific)^5,6^. L1Hs elements encode proteins required to retrotranspose^7^ *in cis*, and to *trans* mobilise *Alu* and SVA retrotransposons and processed mRNAs^8–10^. While the reference genome assembly contains thousands of human-specific TE copies, the vast majority of germline polymorphic TEs found in the global population are non-reference^11,12^. L1Hs-mediated germline insertional mutagenesis is a prominent source of disease, whereas somatic L1Hs retrotransposition can occur during early embryogenesis as well as in the committed neuronal lineage, and is a common feature of many epithelial cancers^13–15^.

A wide array of host factors have been implicated in mammalian TE regulation^16^. Central among them is CpG methylation: most CpGs are located within TEs, and it has been posited that CpG methylation arose to limit the mobility of young TEs^17,18^ whereas older TEs are controlled by repressive histone marks and other pathways. A CpG island present in the L1Hs 5’UTR is usually demethylated as a prerequisite for retrotransposition^19–21^. While TE methylation can be ascertained by locus-specific and genome-wide bisulfite sequencing assays, these approaches are currently limited in throughput and resolution, respectively. Here, we demonstrate the capacity of ONT sequencing to concurrently assess TE methylation and resolve new germline and somatic TE insertions.

We employed an ONT PromethION platform to sequence 5 human samples at ∼15x genome-wide depth. Samples consisted of hippocampus, heart and liver tissue - representing each of the three germ layers - from an individual (CTRL-5413, female, 51yrs) without post-mortem pathology, and paired tumour/non-tumour liver tissue from a second individual (HCC33, female, 57yrs) (Fig. 1a and Supplementary Table 1). ONT analysis allowed us to compare CpG methylation amongst genomes^22^ and between haplotypes within samples^23^. Examining TE subfamilies *en masse*, we observed tumour-specific L1Hs demethylation in HCC33 was more pronounced than demethylation of other young TEs, and of the genome overall (Fig. 1b and Supplementary Fig. 1). Comparing CTRL-5413 normal hippocampus, heart, and liver samples, we found L1Hs methylation decreased in that order (Fig. 1c), an effect that appeared more marked amongst older LINE-1 subfamilies (Supplementary Fig. 2). Older *Alu* subfamilies were also generally less methylated on average than younger elements (Supplementary Fig. 3). SVA methylation, in contrast, did not vary amongst the three samples or with subfamily age (Supplementary Fig. 4). Long terminal repeat (LTR5_Hs) regions flanking the likely immobile human endogenous retrovirus K (HERV-K) family were less methylated than other TEs in normal tissues and non-tumour liver (Fig. 1b,c). Genome-wide and TE subfamily methylation were slightly lower in HCC33 non-tumour liver than CTRL-5413 normal liver (Fig. 1b,c). Composite methylation profiles spanning the previously inaccessible interiors of full-length TEs revealed a clear trough adjacent to the L1Hs 5’UTR CpG island in all samples (Fig. 2), whereas the CpG-rich VNTR (variable number of tandem repeats) core of the youngest SVA_F_ family was more consistently methylated than its flanking SINE-R and *Alu*-like sequences (Fig. 2).

**Fig. 1:**
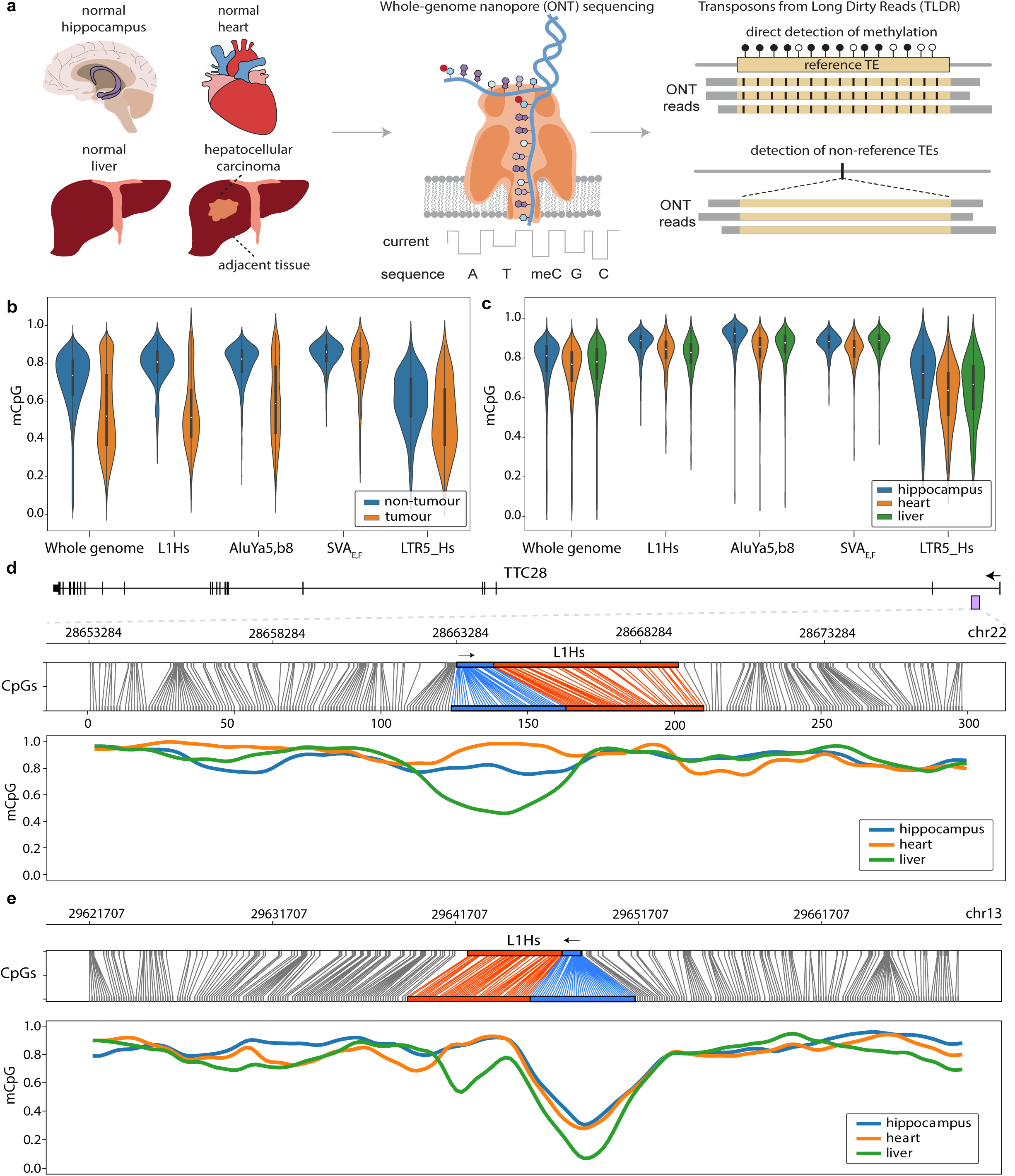
Measurement of CpG methylation on TEs. **(a)** Hippocampus, heart and liver tissue from a healthy individual (CTRL-5413), as well as tumour and adjacent liver tissue from a hepatocellular carcinoma patient (HCC33), were ONT sequenced. TLDR analysis identified TE insertions and quantified TE locus-specific CpG methylation. **(b)** CpG methylation in HCC33 samples for the whole genome (6kbp windows), L1Hs copies >5.9kbp, young *Alu* copies >280bp (AluYa5, AluYb8), human-specific SVA copies >1kbp (SVA_E_, SVA_F_) and HERV-K flanking long terminal repeats >900bp (LTR5_Hs). **(c)** As for (b), except for CTRL-5413 normal tissues. **(d)** Methylation profile of a reference L1Hs intronic to TTC28. A purple rectangle indicates the L1Hs position within the TTC28 locus. Upper panel: relationship between CpG positions in genome space and CpG space. The L1Hs 5’UTR and body are highlighted in blue and orange, respectively. Lower panel: Fraction of methylated CpGs for CTRL-5413 tissues across CpG space. **(e)** Similar to (d), except for an intergenic L1Hs located on chromosome 13 and known to be demethylated and mobile during neurodevelopment^19,27^.

**Fig. 2:**
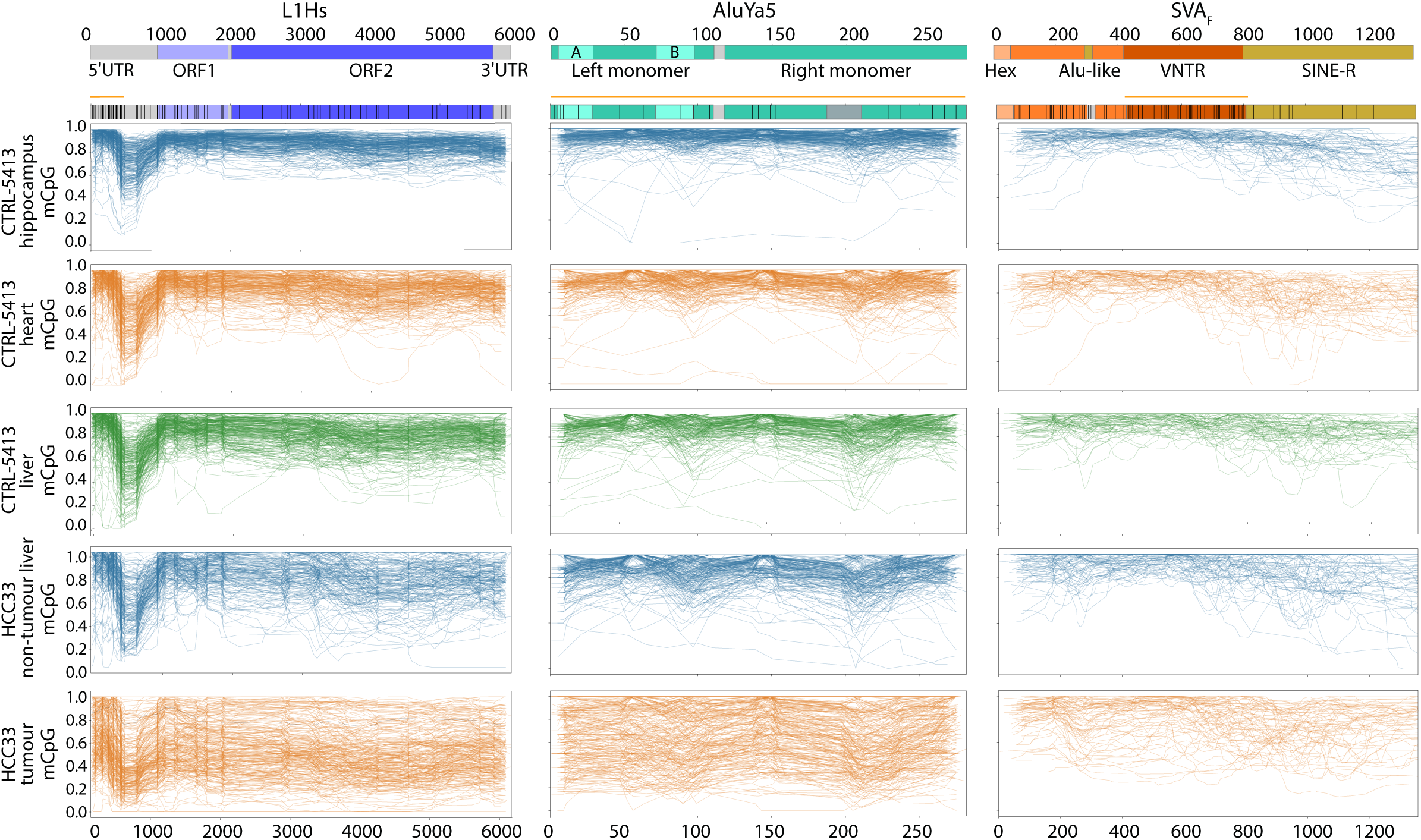
Composite methylation profiles for representative mobile human TE subfamilies. Data are shown for L1Hs, AluYa5 and SVA_F_ in CTRL-5413 and HCC33 samples. Each graph displays up to 300 profiles for the specified TE subfamily. Annotated TE consensus sequences are provided at top, with CpG positions (black bars) and CpG islands (orange lines) indicated.

Whilst most TEs are constitutively methylated (Supplementary Fig. 5a) we identified striking patterns of differential methylation for individual reference genome TEs (Supplementary Fig. 5b-d, Supplementary Table 2). For example, an L1Hs located intronic to the TTC28 gene and known to be mobile in liver and other cancers^24–26^ was hypomethylated in CTRL-5413 liver (Fig. 1d). A slightly 5’ truncated L1Hs situated on chromosome 13 and found, thus far, to cause somatic retrotransposition during neurodevelopment in two unrelated individuals^19,27^ was strongly demethylated in each CTRL-5413 sample (Fig. 1e). An L1Hs situated antisense and intronic to ZNF638 was similarly demethylated, particularly in CTRL-5413 heart tissue, and from its 5’UTR promoted a previously described alternative ZNF638 transcript^28^ (Supplementary Fig. 5c). A chromosome 1 L1Hs that is mobile in the germline and cancer^29^, and recently highlighted as expressed in senescent fibroblasts^30^, was hypomethylated in CTRL-5413 heart and liver, but not hippocampus (Supplementary Fig. 5d). Despite an overall trend towards tumour-specific L1Hs demethylation in HCC33, we also noted exceptional TEs that were hypermethylated in tumour relative to non-tumour liver, such as an L1Hs copy intronic to PGAP1 (Supplementary Fig. 6). As well as examples of TEs methylated exclusive of the surrounding locus, we found TEs apparently demethylated by virtue of their genomic location (Supplementary Fig. 7). By generating read-backed phased methylation profiles^23,31,32^ we found haplotype-specific differentially methylated regions within imprinted genes, such as GNAS and PEG3 (Supplementary Fig. 8), and in individual reference TE copies (Supplementary Fig. 9). These results highlight how haplotype-specific TE regulation can be studied - and placed amid a wider genomic context - via ONT analysis.

To study non-reference TE insertions, we developed TLDR (Transposons from Long Dirty Reads), a software tool that detects, assembles and annotates insertions via long-read alignments, including ONT and PacBio sequencing data. A major feature of TLDR is that it can resolve entire TE insertions, along with transductions, 5’ inversions, target site duplications (TSDs), 3’ poly(A) tracts and other hallmarks of LINE-1 mediated retrotransposition^1^ (Fig. 3a, Supplementary Fig. 10, and Supplementary Table 3) while achieving sensitivity similar to our short-read TE insertion detection method, TEBreak. Highlighting its capacity to detect somatic retrotransposition events, TLDR successfully re-identified both PCR-validated tumour-specific L1Hs insertions previously found by us with short-read sequencing of patient HCC33 samples^33^. TLDR accurately recapitulated their LINE-1 insertion features, including TSDs, and now revealed the internal breakpoint of the ∼2kb 5’ inversion of the EFHD1 insertion (Fig. 3a and Supplementary Table 3). No additional HCC33 tumour-specific TE insertions were found by TLDR.

**Fig. 3:**
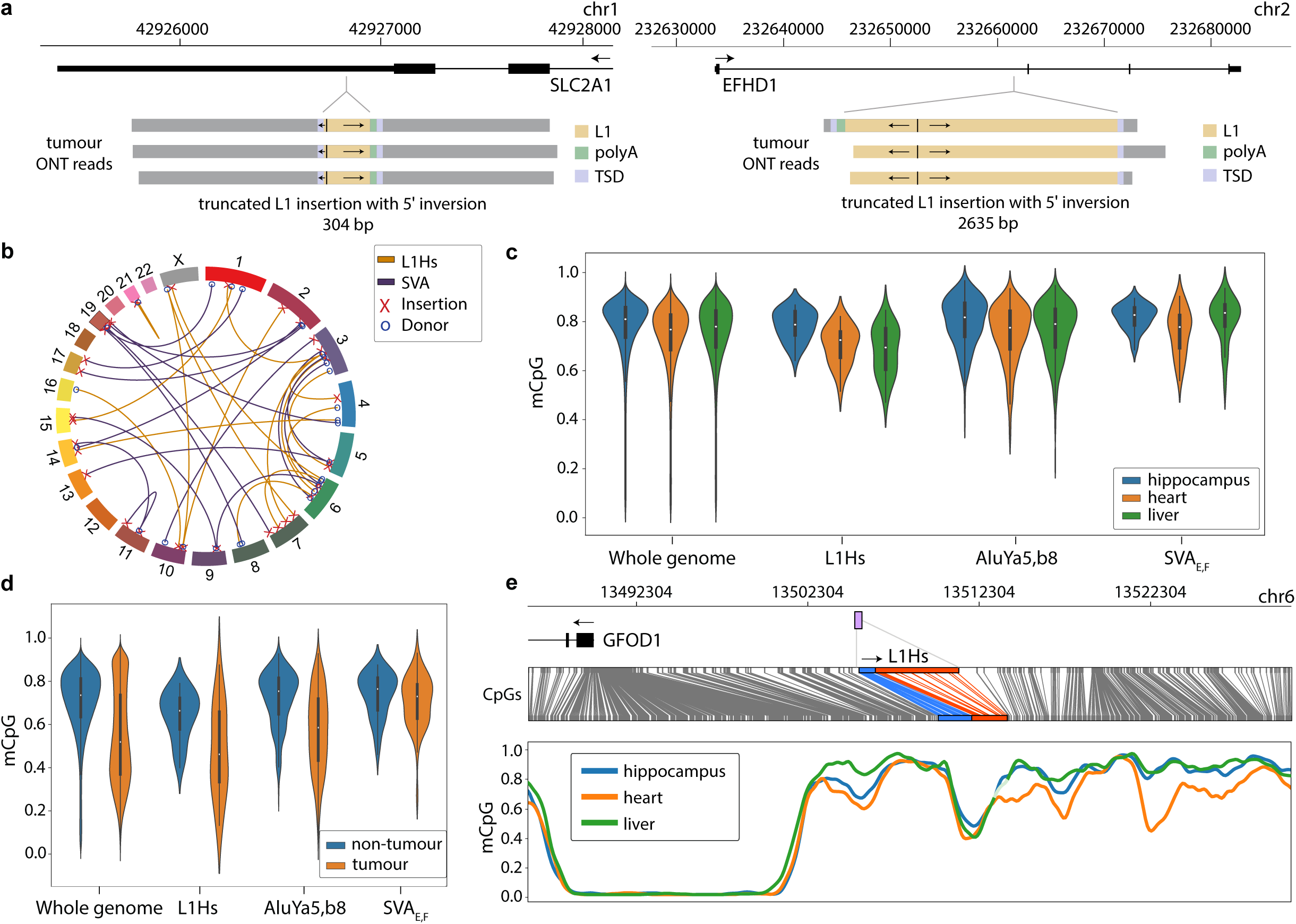
Non-reference TE detection and assessment of CpG methylation. **(a)** Detection of HCC33 tumour-specific 5’ truncated L1Hs insertions. Arrows within L1Hs sequences indicate 5’ inversions. **(b)** Source element (donor) to insertion relationships for germline LINE-1 and SVA insertions identified in CTRL-5413 and HCC33 by TLDR. **(c)** Fraction of methylated CpGs in CTRL-5413 tissues, as in Fig. 1c, except for non-reference TE insertions. **(d)** As for (c), except for non-reference TE methylation in HCC33 samples. **(e)** Exemplar methylation profile for a non-reference L1Hs (UUID d11b3baf, Supplementary Table 3a). A purple box indicates the genomic position of the L1Hs upstream of GFOD1 on chromosome 6. The liver smoothed plot line is coloured to appear faded for a short lower confidence region (<20 methylated/demethylated calls within a 30 CpG window). Panels are otherwise as described for Fig. 1d,e. Note: the demethylated region to the left of the L1Hs sequence corresponds to the GFOD1 promoter.

To assess the sensitivity of TLDR for germline TE polymorphisms, we compiled a high-confidence set of known non-reference (KNR) TE insertions reported by at least two previous studies (Online Methods). We then applied TEBreak to ∼45x Illumina whole genome sequencing generated from CTRL-5413 heart and HCC33 non-tumour liver. Of the high-confidence germline KNR set, 2464 were detected by TLDR in CTRL-5413 or HCC33 (Supplementary Table 3) and 2533 were called by TEBreak (Supplementary Table 6). A total of 2357 KNR insertions were reported by both TLDR and TEBreak (Jaccard metric ≅0.96), indicating high concordance. Consistent with the recent use of PacBio long-read sequencing to resolve LINE-1 insertions in difficult to map genomic regions^34^, we found non-reference insertions called only by TLDR covered a much broader spectrum of mappability scores than those found only by TEBreak or both methods (Supplementary Fig. 11). Notably, TLDR reports useful sequence features of non-reference TE insertions^1^. These include, for example, 5’ and 3’ transductions carried by LINE-1 and SVA insertions^35^, of lengths ranging here from 31bp to 2072bp, and attributable to their putative source elements (Fig. 3b and Supplementary Table 4). Internal polymorphisms, such as VNTR length differences within SVAs (Supplementary Table 3d) are also resolved by TLDR.

While somatic methylation appears less ubiquitous as TE subfamilies age (Supplementary Figs. 2-4), the initial duration required for new TE insertions to be strictly methylated is unclear. Using TLDR, we established that non-reference TE insertions (Fig. 3c,d) appeared to be, on average, less methylated than reference elements (Fig. 1b,c) in each of the CTRL-5413 tissues and HCC33 non-tumour liver. Reference (82.5%) and non-reference (70.6%) L1Hs insertions exhibited the largest difference (P<5.76e-30, Mann-Whitney test) amongst the analysed TE subfamilies. Of 59 germline TE insertions (10 L1Hs, 46 AluY, 3 SVA) detected by both TLDR and TEBreak in either CTRL-5413 or HCC33, and absent from prior studies considered here (Online Methods), we identified 4 full-length L1Hs instances with an intact 5’UTR CpG island. The average methylation observed for these 4 potentially recent L1Hs insertions was 65.4%, trending less than the overall non-reference L1 cohort (P<0.056, Mann-Whitney test). As for individual reference TEs, TLDR can be used to generate element-specific methylation profiles for non-reference TE insertions (Fig. 3e, Supplementary Fig. 12 and Supplementary Table 5), including retrotransposition-competent source L1Hs copies. For example, the non-reference element L1.2, responsible for the first report of LINE-1 mobility and pathogenesis in humans^7,36^, was ∼15% less methylated in CTRL-5413 liver and heart than in hippocampus (Supplementary Fig. 12a, Supplementary Table 5). As the vast majority of mobile L1Hs copies in the global population are absent from the reference genome^6^, the capacity of TLDR to find non-reference L1Hs alleles and survey their methylation state in parallel is notable.

Interrogated with TLDR, long-read ONT sequencing has unprecedented utility to detect and characterise somatic and germline TE insertions, including in genomic regions refractory to reliable short-read mapping. ONT analysis can provide end-to-end resolution of TE insertions, without generating molecular artifacts associated with PCR amplification. Hallmark features of LINE-1 mediated retrotransposition are therefore readily recovered by TLDR, including relatively long transductions and internal rearrangements. While bisulfite sequencing depicts CpG methylation at TE termini, ONT analysis yields a direct methylation readout throughout TE sequences. These attributes mean TLDR has the potential to, for instance, resolve somatic TE insertions arising during neurodevelopment, and at the same time infer methylation of the inserted TE, as well as its integration site and source locus. As shown here for the SVA VNTR and 3’ end of the L1Hs 5’UTR, CpG methylation may vary greatly within mobile TEs. These “sloping shores” of methylation around recently inserted TE CpG islands^37^ have the potential to mislead assays that mainly access TE termini. While L1Hs locus-specific bisulfite sequencing^19^ generates data concordant with those shown here, ONT analysis is far higher throughput and encompasses all human TE subfamilies. TLDR therefore makes a broad range of new questions relating to TE biology more accessible.

## Acknowledgements

The authors thank the human subjects of this study who donated tissues to the MRC Edinburgh Brain and Tissue Bank and the Centre Hépatobiliaire, Paul-Brousse Hospital. The authors thank P. Gerdes for helpful discussions, and acknowledge the Translational Research Institute (TRI) for research space, equipment and core facilities that enabled this research. This study was funded by the Australian Department of Health Medical Frontiers Future Fund (MRFF) (MRF1175457 to ADE), the Australian National Health and Medical Research Council (NHMRC) (GNT1125645, GNT1138795, GNT1173711 to GJF), an NHMRC Early Career Fellowship (GNT1161832) to SWC, a CSL Centenary Fellowship to GJF, and by the Mater Foundation (Equity Trustees / AE Hingeley Trust).

ADE, SWC, and GJF designed the research project, and ADE, NS, SWC and GJF wrote the manuscript. SWC conducted sample preparation and quality control. FJS-L, JF and PMB provided resources. ADE and NS developed software tools and analysed the data.

## Methods

See Online Methods.

