## Supplementary Figures and Methods for "Nanopore sequencing enables comprehensive transposable element epigenomic profiling"

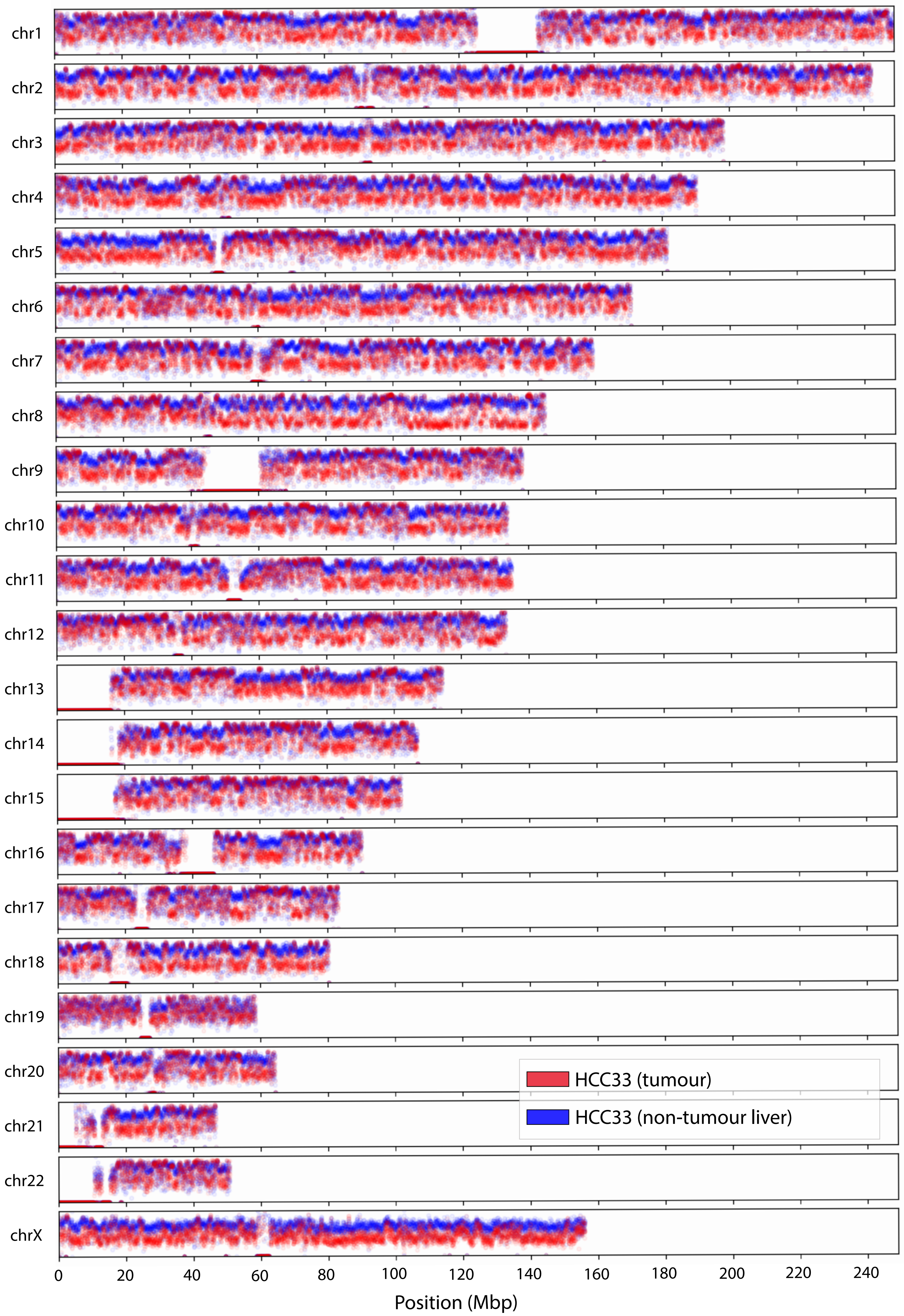


**Supplementary Fig. 1: Whole genome ONT methylation profiles for patient HCC33 samples.** Comparison of liver tumour (red) and non-tumour liver (blue) tissues indicates genome-wide demethylation of the tumour genome. The reference genome is binned into 25kbp segments.

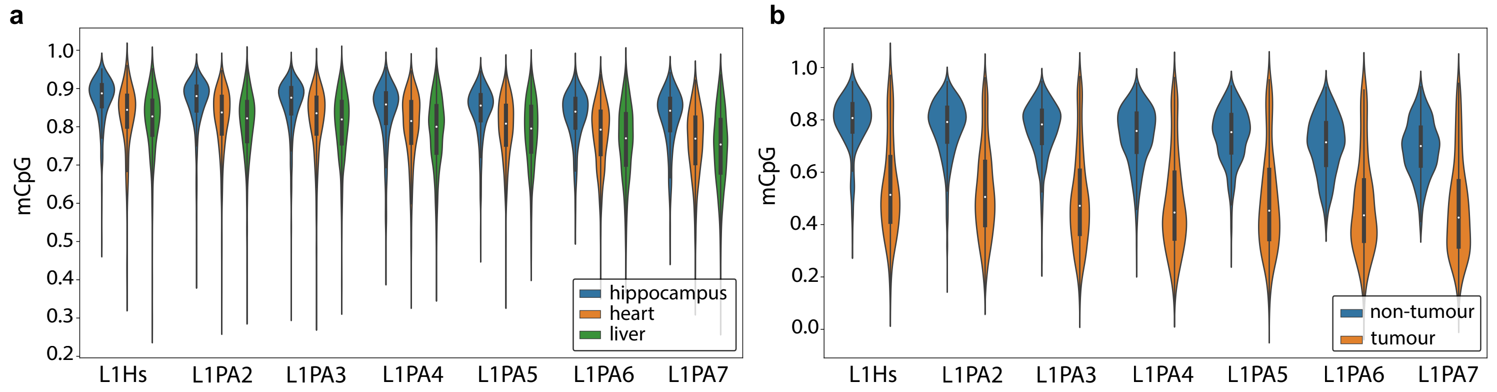


**Supplementary Fig. 2: Violin plots depicting methylated CpG fractions for LINE-1 subfamilies.** L1Hs is still mobile and is the only human-specific LINE-1 subfamily, followed in increasing age by L1PA2 through L1PA7, which are shared with other great ape species and are immobile in humans. At least 10 methylation calls (+ or -) were required in each sample for an element to be included. Results are shown for **(a)** CTRL-5413 normal hippocampus, heart, and liver tissues and **(b)** patient HCC33 tumour and non-tumour liver samples.

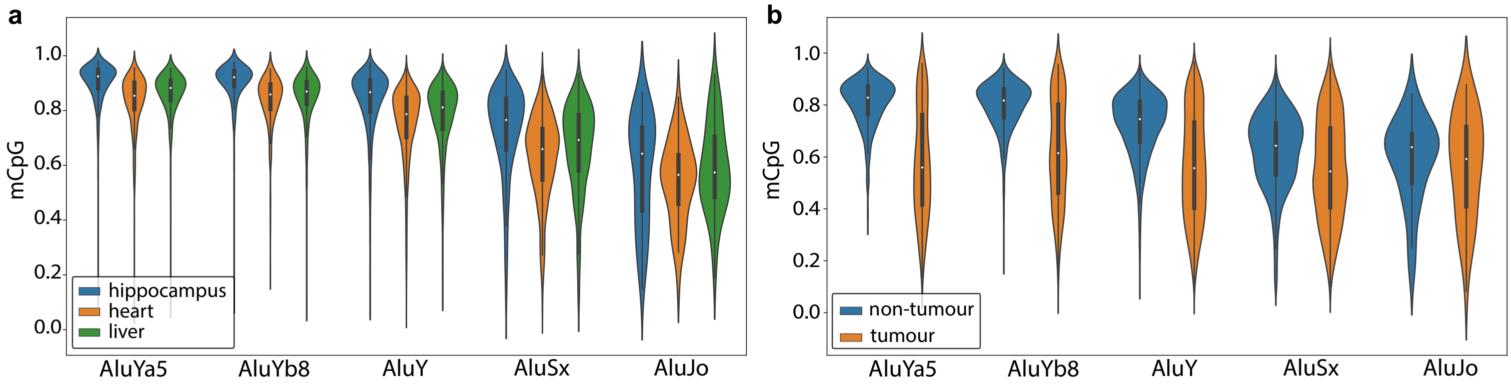


**Supplementary Fig. 3: Violin plots depicting methylated CpG fractions for *Alu* subfamilies.** AluYa5 and AluYb8 are the most active *Alu* subfamilies in the human genome. Other high-copy number *Alu* subfamilies are shown for comparison. AluSx and AluJo elements are generally older than AluY elements. At least 10 methylation calls (+ or -) were required in each sample for an element to be included. Results are shown for **(a)** CTRL-5413 normal hippocampus, heart, and liver tissues and **(b)** patient HCC33 tumour and non-tumour liver.

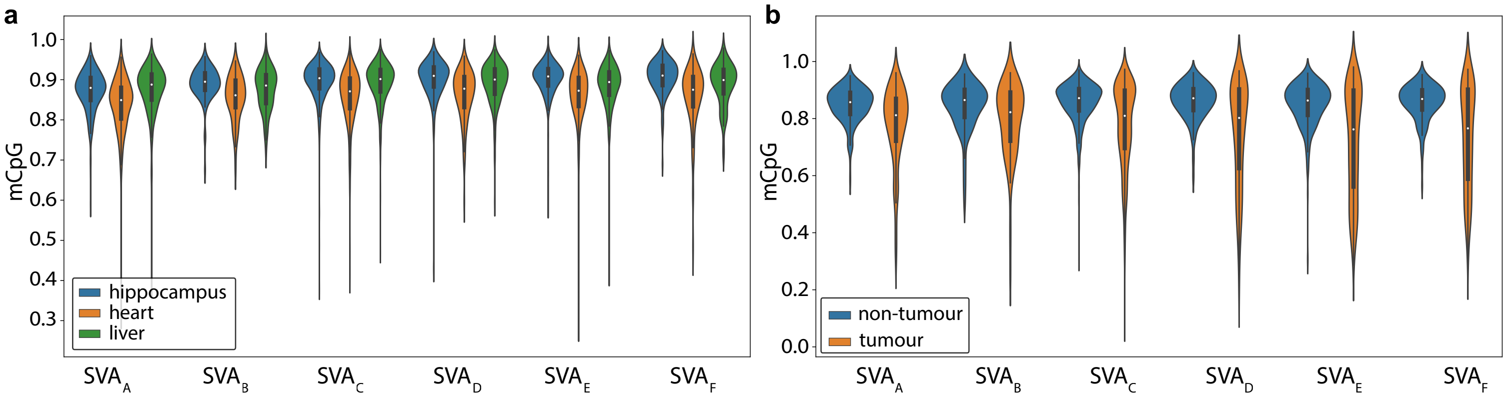


**Supplementary Fig. 4: Violin plots depicting methylated CpG fractions for SVA subfamilies.** $SVA_F$ is the most recently emerged subfamily, followed by $SVA_E$ through $SVA_A$ in increasing age. At least 10 methylation calls (+ or -) were required in each sample for an element to be included. Results are shown for **(a)** CTRL-5413 normal hippocampus, heart, and liver tissues and **(b)** patient HCC33 tumour and non-tumour liver.

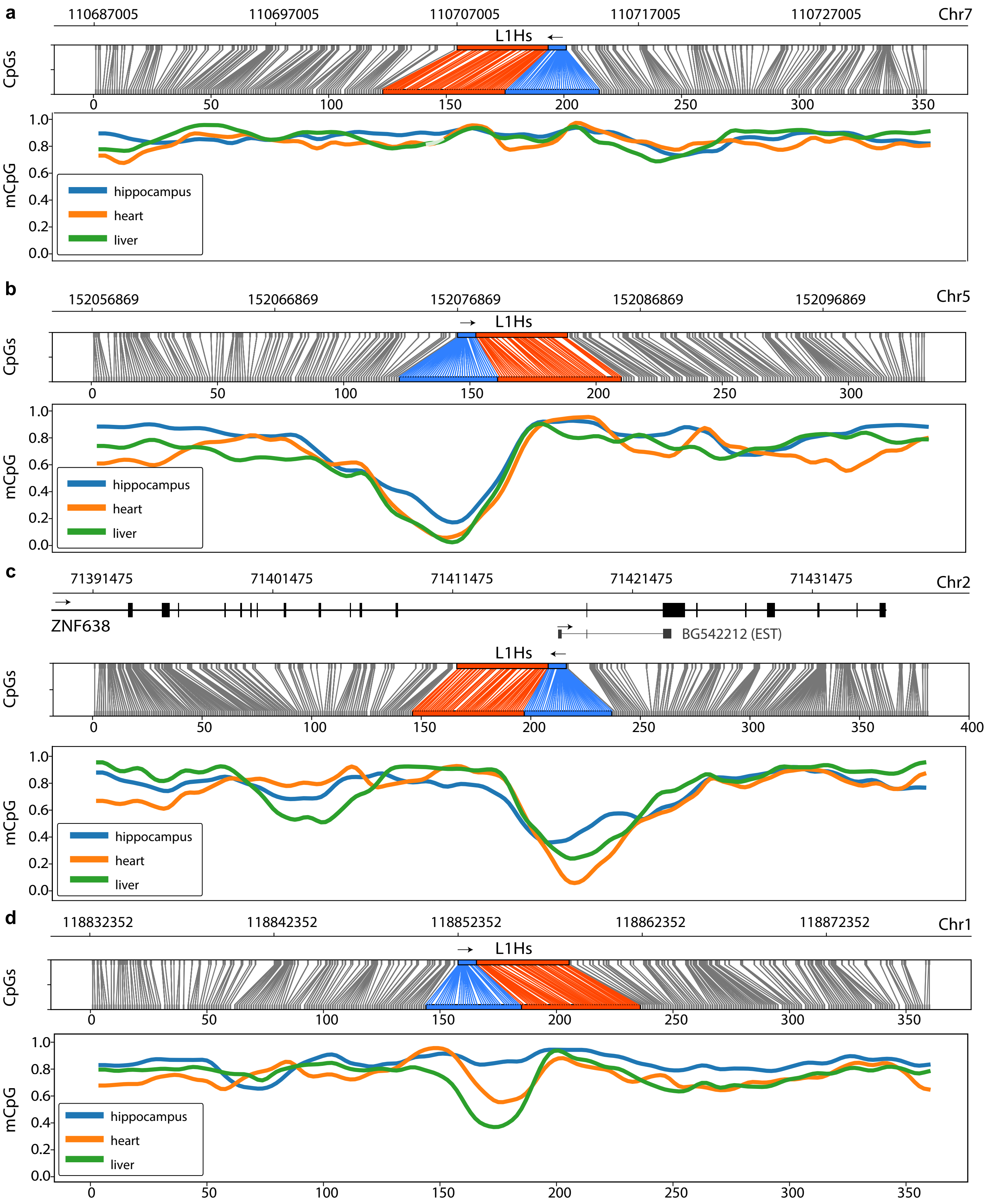
a
110687005
110697005
110707005
110717005
110727005
Chr7
L1Hs
CpGs
mCpG
hippocampus
heart
liver
b
152056869
152066869
152076869
152086869
152096869
Chr5
L1Hs
CpGs
mCpG
hippocampus
heart
liver
c
71391475
71401475
71411475
71421475
71431475
Chr2
ZNF638
BG542212 (EST)
L1Hs
CpGs
mCpG
hippocampus
heart
liver
d
118832352
118842352
118852352
118862352
118872352
Chr1
L1Hs
CpGs
mCpG
hippocampus
heart
liver

**Supplementary Fig. 5: Reference L1Hs methylation profiles. (a)** A broadly methylated intergenic L1Hs located on chromosome 7. Upper panel: correspondence between CpG positions in genome space and CpG space. The L1Hs 5′UTR and body are highlighted in blue and orange, respectively. Lower panel: fraction of methylated CpGs for CTRL-5413 tissues across CpG space. Data are shown via a sliding window plot. **(b)** As for (a), except showing a strongly demethylated element on chromosome 5. **(c)** As for (a), except for an element intronic to ZNF638 and hypomethylated in each tissue, particularly heart. Transcription of an alternative isoform of ZNF638 from the antisense promoter of this element is supported by an expressed tag sequence (EST). **(d)** As for (a), except showing an element located on chromosome 1 and hypomethylated only in heart and liver. This element was reported as expressed in senescent fibroblasts by de Cecco et al.[18] Note: elements shown in panels (a) and (b) correspond to Chr7$\Delta 12_{L1}$ and Chr5$\Delta 31_{L1}$, respectively, from Sanchez-Luque et al.[19] Each element was methylated consistent with results obtained from this prior study using locus-specific Illumina bisulfite sequencing. The heart sample smoothed plot line in panel (a) is coloured to appear faded for a short lower confidence region (<20 methylated/demethylated calls within a 30 CpG window).

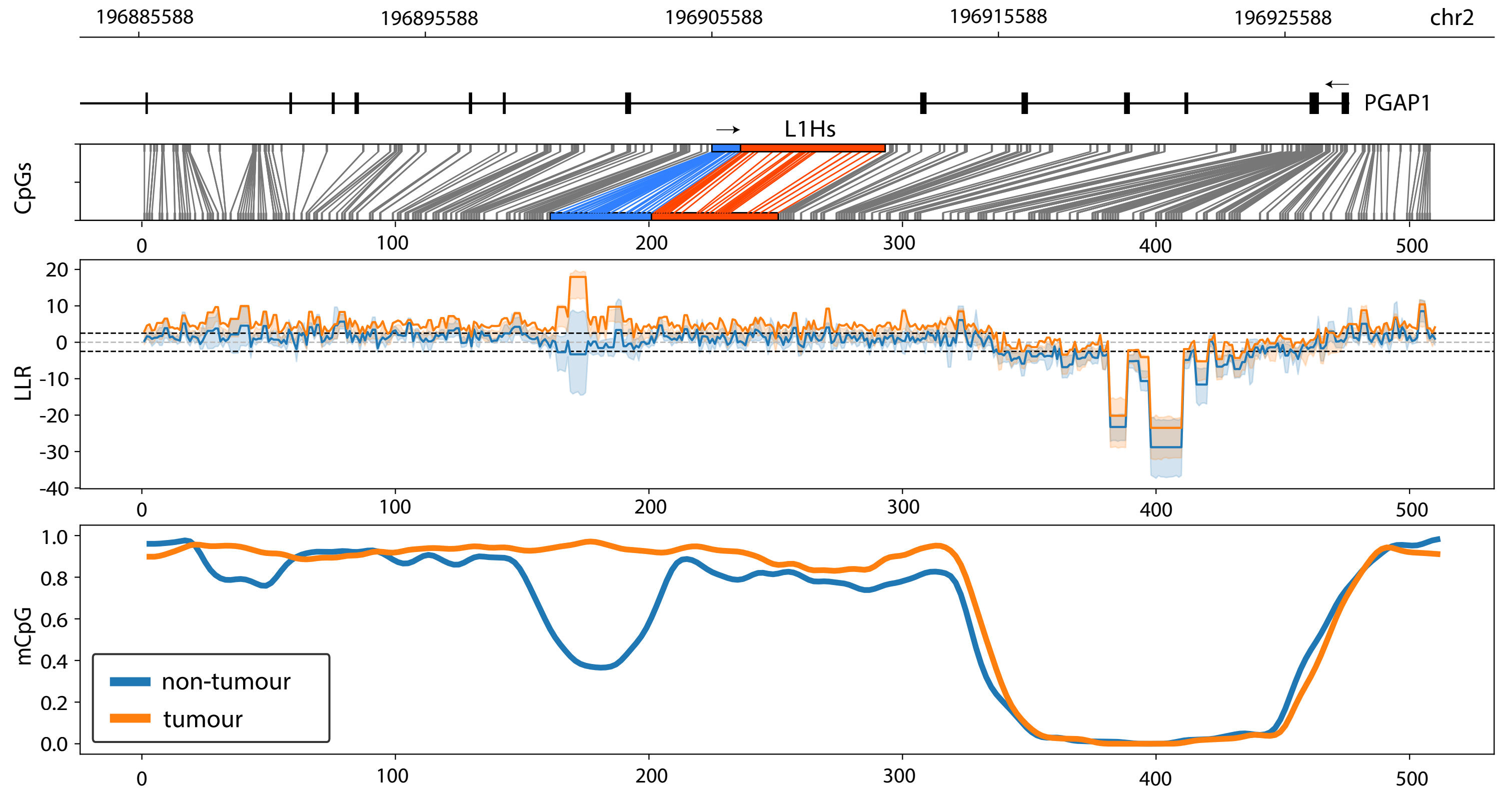


**Supplementary Fig. 6: A reference L1Hs insertion demethylated distinct to the surrounding locus in patient HCC33 non-tumour liver, relative to the matched tumour.** From top to bottom, this figure shows i) the genomic position of the L1Hs in an intron of the PGAP1 gene on chromosome 2, including 20kbp up and downstream of the L1Hs, ii) a diagram showing the correspondence between genome space and CpG space, iii) the log-likelihood ratios (LLRs) for methylation across the region in HCC33 tumour (orange) and non-tumour liver (blue) samples, and iv) the fraction of methylated CpGs for each HCC33 tissue in CpG space. Data are shown via a sliding window plot. The demethylated region to the right of the L1Hs sequence corresponds to the PGAP1 promoter region.

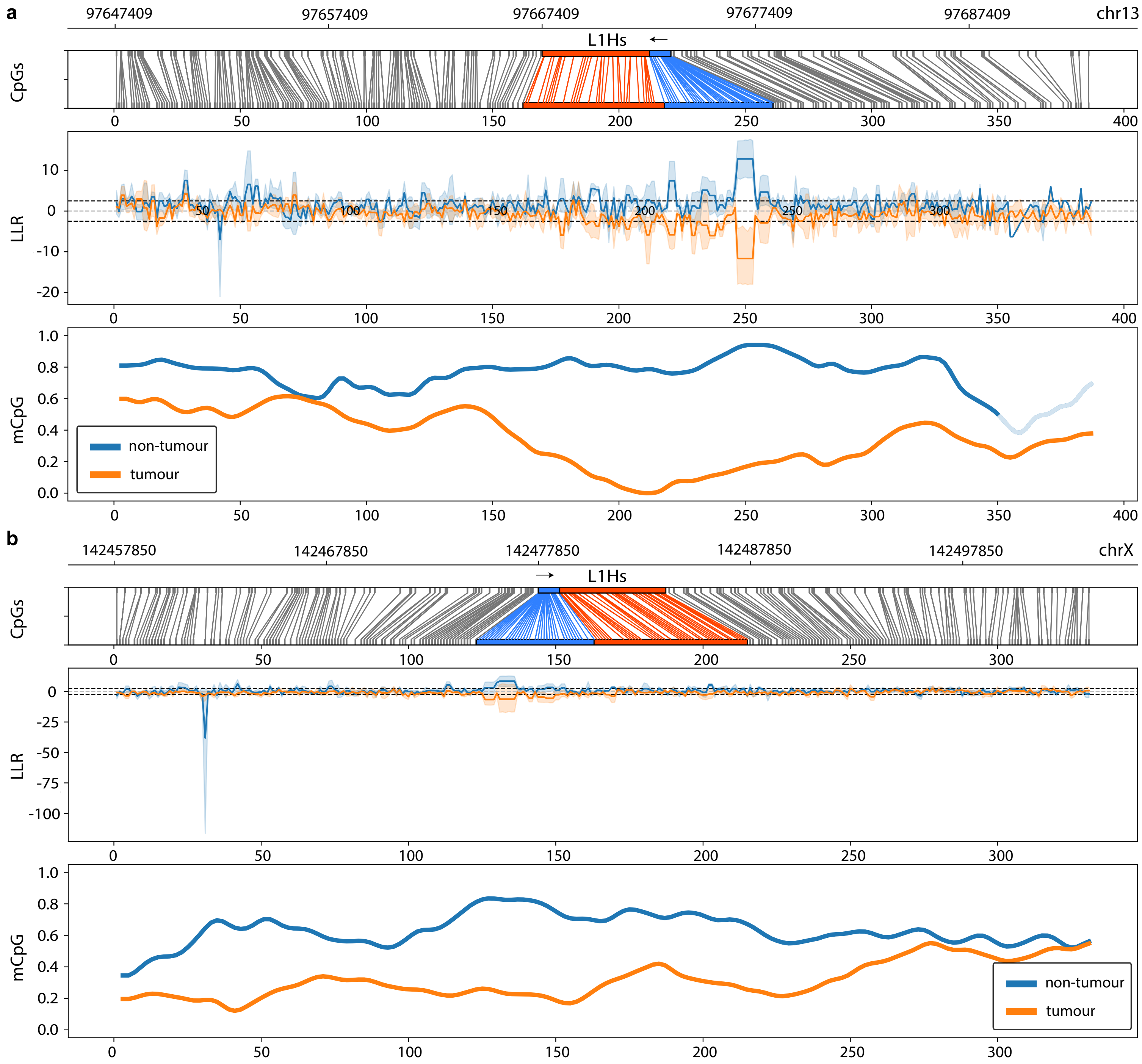


**Supplementary Fig. 7: Example reference L1Hs insertions demethylated along with their surrounding genomic region in patient HCC33 tumour. (a)** From top to bottom, this figure shows i) the genomic position of an intergenic L1Hs on chromosome 12, ii) a diagram showing the correspondence between genome space and CpG space, iii) the log-likelihood ratios (LLRs) for methylation across the region in HCC33 tumour (orange) and non-tumour liver (blue) samples, and iv) the fraction of methylated CpGs for each HCC33 tissue in CpG space. Data are shown via a sliding window plot. **(b)** As for (a), except for an L1Hs located on chromosome X. Note: the non-tumour sample smoothed plot line in panel (a) is coloured to appear faded for a short lower confidence region (<20 methylated/demethylated calls within a 30 CpG window).

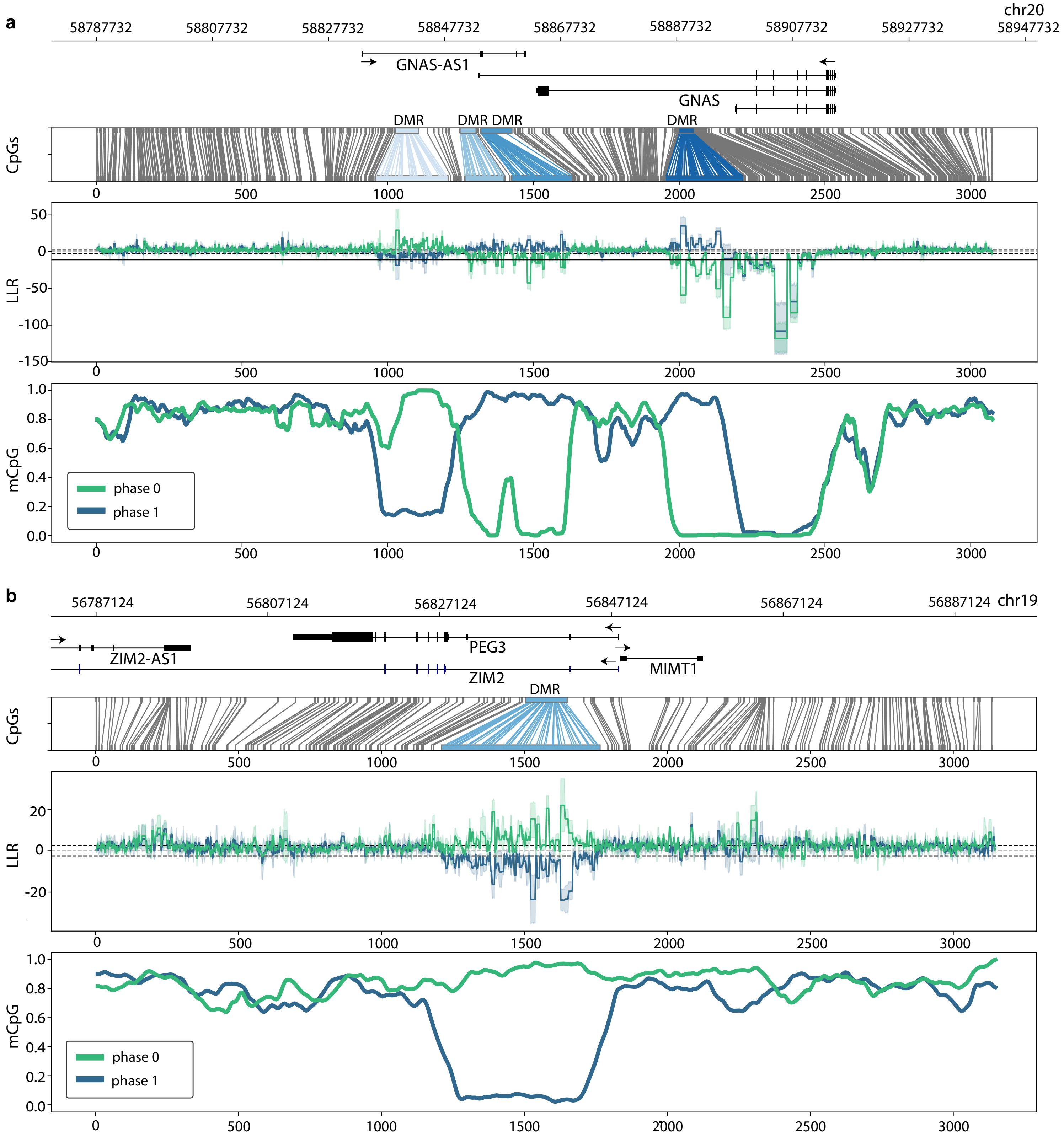


**Supplementary Fig. 8: Known imprinted differentially methylated regions (DMRs) detected via ONT analysis.** Read-backed phasing was used to identify haplotype-specific differences in CTRL-5413 hippocampus methylation for **(a)** the GNAS gene, where DMRs specific to the paternal and maternal alleles were identified, and **(b)** the PEG3 gene, which is known to be expressed from only the paternal allele. Each panel shows, from top to bottom, i) the genomic position of DMRs, ii) a diagram showing the relationship between genome space and CpG space, iii) the log-likelihood ratios (LLRs) for methylation across the region where one haplotype (phase 0, teal) is compared to the other (phase 1, blue), and iv) the fraction of methylated CpGs for each phase. Data are shown via a sliding window plot.

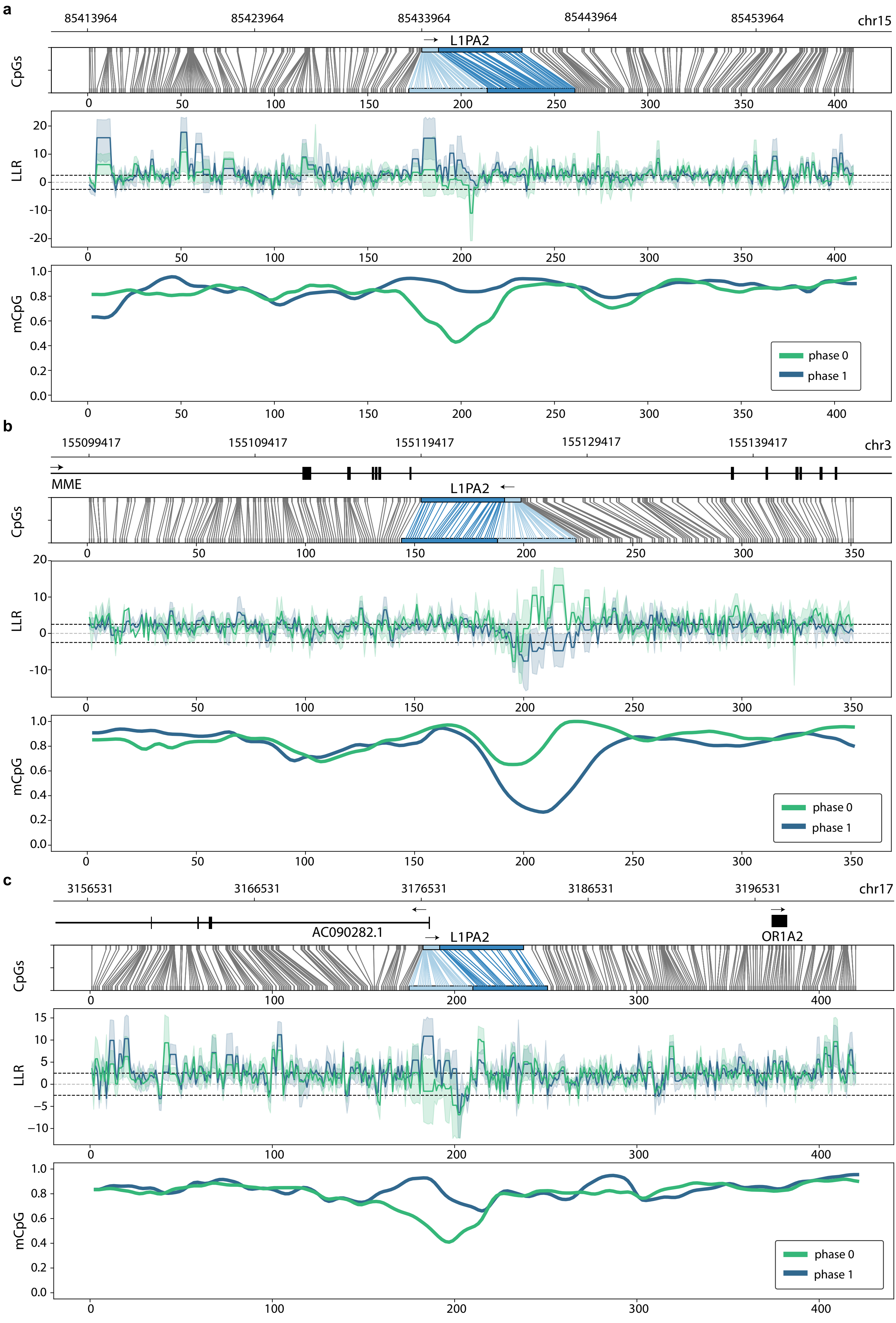

a
85413964
85423964
85433964
85443964
85453964
chr15
L1PA2
CpGs
LLR
mCpG
phase 0
phase 1
b
155099417
155109417
155119417
155129417
155139417
chr3
MME
L1PA2
CpGs
LLR
mCpG
phase 0
phase 1
c
3156531
3166531
3176531
3186531
3196531
chr17
AC090282.1
L1PA2
OR1A2
CpGs
LLR
mCpG
phase 0
phase 1

**Supplementary Fig. 9: Examples of TEs with evidence for haplotype-specific differential methylation.** Read-backed phasing was used to identify haplotype-specific differences in CTRL-5413 hippocampus methylation. Elements in panels **(a-c)** are full-length members of the L1PA2 subfamily. Each panel shows, from top to bottom, i) the genomic position of an L1PA2, ii) a diagram showing the relationship between genome space and CpG space, iii) the log-likelihood ratios (LLRs) for methylation across the region where one haplotype (phase 0, teal) is compared to the other (phase 1, blue), and iv) the fraction of methylated CpGs for each phase. Data are shown via a sliding window plot. The 5′UTR and body of each L1PA2 element are highlighted in light blue and blue, respectively.

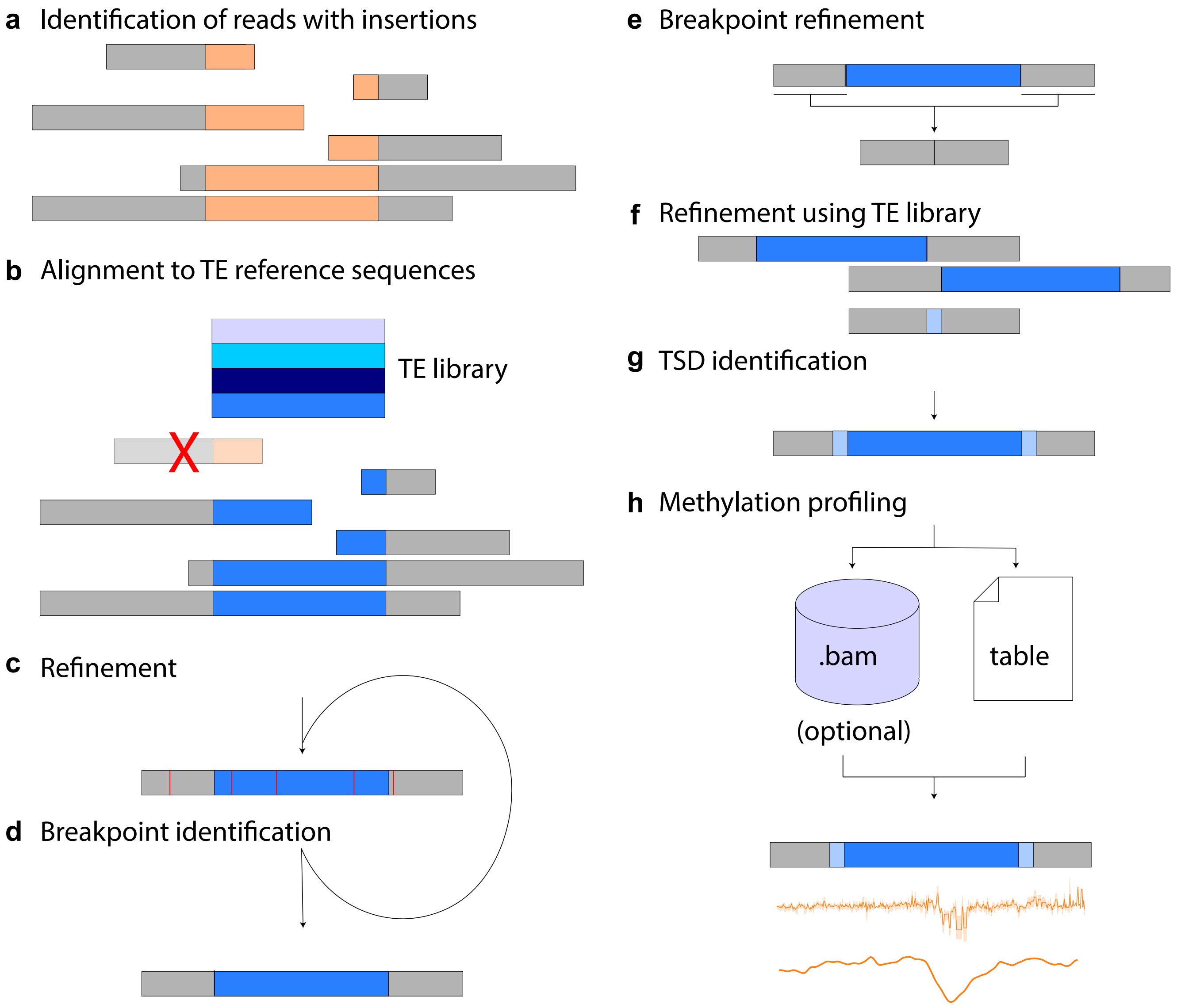


**Supplementary Fig. 10: Diagram depicting the operation of TLDR.** Starting from a cluster of reads **(a)** with a profile of mapped (grey) and unmapped (orange) regions consistent with an insertion, the unmapped portion of reads is aligned to a TE reference library **(b)**, and a consensus is built from multiple sequence alignment of the reads supporting a TE insertion **(c)**, which is refined through realignment and pileup-based assessment of the supporting reads **(d)**. The TE breakpoints are initially determined approximately and are used to select the corresponding segment of the reference genome **(e)**. The reference is then realigned to the refined consensus to better define the TE breakpoints **(f)**. Subsequent refinement using alignments to the TE reference library define target site duplications (TSDs) if present **(g)**, and assist with final annotation of the insertion. An optional per-insertion .bam file is generated which, along with the consensus sequence and flanking regions, can be used to determine the methylation status of the insertion **(h)**. For further details please see Online Methods.

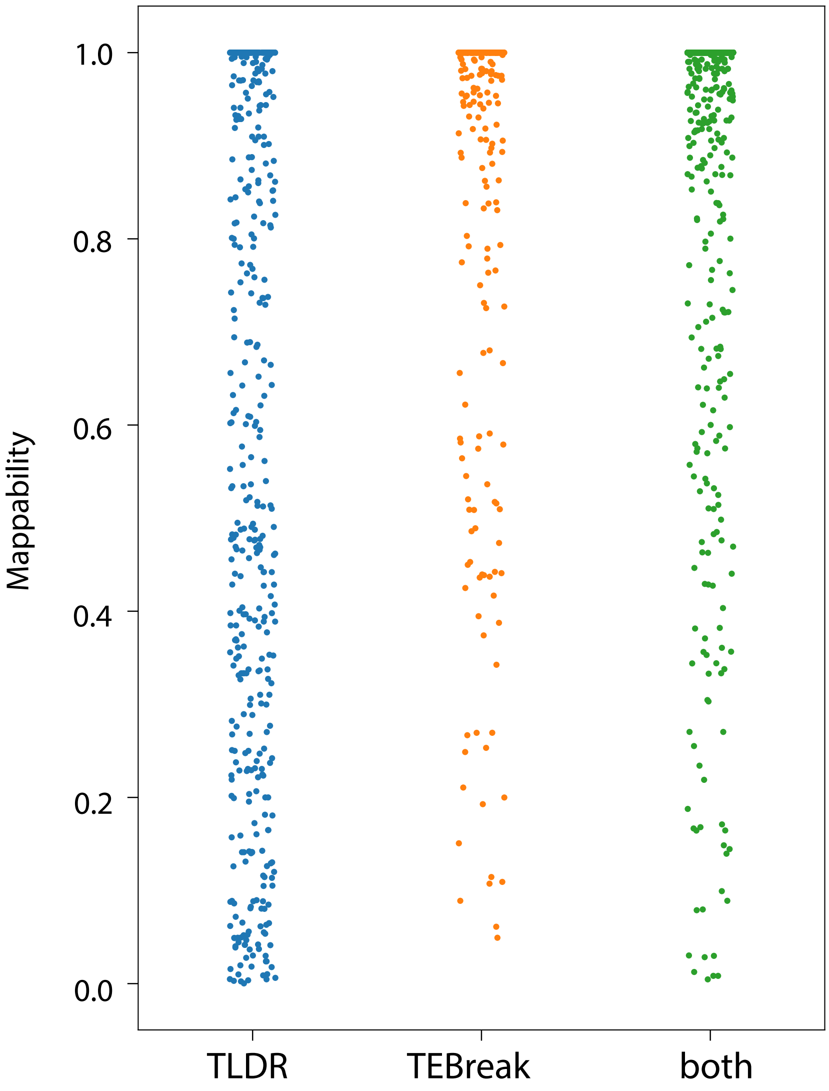


**Supplementary Fig. 11: Insertions called by TLDR cover a broad spectrum of mappability.** Mappability was calculated for a 200bp window centered on each insertion, with insertions categorised as detected by TLDR only, TEBreak only, and overlapping calls of TLDR and TEBreak (“both”). Only insertion calls passing all filters were considered. Mappability was determined via the GEM mappability method. Please see Online Methods for further details.

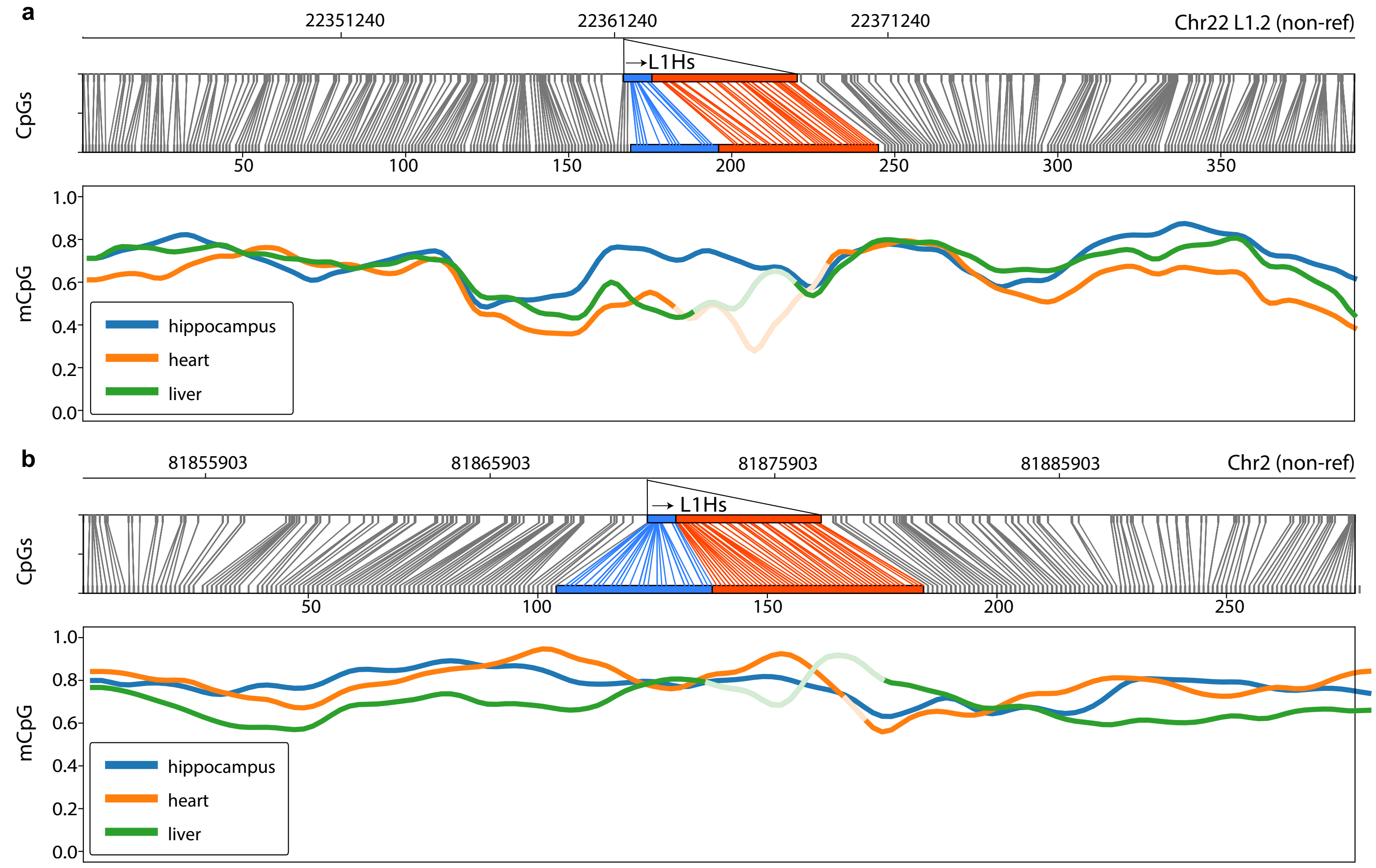


**Supplementary Fig. 12: Non-reference L1Hs methylation profiles. (a)** The 5′UTR of a mobile L1Hs (L1.2)[20] located on chromosome 22 is less methylated in CTRL-5413 heart and liver than hippocampus. Upper panel: correspondence between CpG positions in genome space and CpG space. The L1Hs 5′UTR and body are highlighted in blue and orange, respectively. Lower panel: fraction of methylated CpGs for CTRL-5413 tissues across CpG space. Data are shown via a sliding window plot. **(b)** As for (a), except showing a broadly methylated element on chromosome 12. This L1Hs corresponds to Chr2Δ2L1 from Sanchez-Luque et al.[19] and was methylated consistent with results obtained from this prior study using locus-specific Illumina bisulfite sequencing. Note: smoothed plot lines are coloured to appear faded for lower confidence regions (<20 methylated/demethylated calls within a 30 CpG window).

### Online Methods

#### Human tissue samples

Snap frozen hippocampus, heart and liver tissue from one post-mortem individual (CTRL-5413) without neurological disease was provided by the Edinburgh Sudden Death Brain and Tissue Bank with ethical approval to be used as described in the study (East of Scotland Research Ethics Service, Reference: LR/11/ES/0022). Further ethics approvals were provided by the Mater Health Services Human Research Ethics Committee (Reference: HREC-15-MHS-52) and the University of Queensland Medical Research Review Committee (Reference: 2014000221). Liver tumour and non-tumour samples were previously obtained from a patient (HCC33) who underwent surgical resection at the Centre Hépatobiliaire, Paul-Brousse Hospital and were analysed with approval from the French Institute of Medical Research and Health (Reference: 11-047).

#### Sequencing data generation

Tissues were subjected to phenol-chloroform DNA extraction and quantified on an Agilent Tapestation. DNA libraries were prepared at the Australian Genome Research Facility (AGRF) using the genomic DNA by ligation kit (SQK-LSK109) and were sequenced on an Oxford Nanopore Technologies (ONT) PromethION platform (r9.4.1 chemistry). Yield and read N50s varied amongst samples (Supplementary Table 1). Bases were called using guppy version 1.8.5 (Oxford Nanopore Technologies) and aligned to the reference genome build hg38 using minimap2 version 2.16[1] and samtools version 1.9[2]. Reads were indexed and per-CpG methylation calls generated using nanopolish version 0.11.0[3]. Methylation likelihood data were sorted by position and indexed using tabix version 1.9[4].

To generate short reads for comparison of TE insertion detection methods and for the generation of high-quality variant calls necessary for haplotyping, DNA for one sample from each individual (HCC33 non-tumour and CTRL-5413 heart), was sequenced to 45x depth by AGRF on an Illumina NovaSeq 6000. Reads were aligned to hg38 using bwa mem version 0.7.12[5] and samtools version 1.9 and duplicate reads were marked using picard tools version 2.18.0[6]. Variants were called using GATK Haplotype Caller version 3.7[7] and known variants annotated via SnpSift version 4.3t[8] using dbSNP build 146[9]. Read-backed phasing of ONT reads was done using whatshap version 0.18[10].

#### Data Availability

ONT and Illumina sequencing data were deposited in the Sequence Read Archive (SRA) using the BioProject identifier PRJNA629858.

#### Reference insertions

Per-element methylation statistics for reference TEs were generated using a python script, segmeth.py, available in the te-nanopore-tools github repository at https://github.com/adamewing/te-nanopore-tools. Reads mapping completely within TEs were excluded from the reference TE methylation analysis to minimise the possibility of mismapping. Reference TE locations were derived from the repeatmasker[11] .out files available for hg38 from the UCSC Genome Browser[12]. As SVA elements are often broken into multiple adjacent SVA annotations[13], we merged adjacent similarly oriented SVAs prior to analysis and considered elements annotated as longer than 1000bp. LINE-1 elements were considered if annotated as greater than 5900bp in length. Only *Alu* elements greater than 280bp were considered. We required at least 5 methylation calls i.e. abs(log-likelihood ratio) > 2.5 across all samples to include an element in the survey (Supplementary Table 2).

Methylation plots for individual elements can be generated using the plotmeth_ref_multi.py or plotmeth_ref_hap.py scripts; the former generates plots for one or more samples and the latter generates plots for .bams that have been haplotype tagged using whatshap. These plotting scripts generate plots with four panels: a positional panel which includes a rudimentary depiction of gene models, a panel which shows the conversion from genome space (i.e. A,C,T,G) into CpG space while optionally highlighting one or more segments, a plot of log-likelihood ratios via seaborn[14], and a plot showing the fraction of methylated CpGs which is windowed and stepped according to user parameters and smoothed using a Hann function. Methylation data displayed here were plotted using a 30bp sliding window with a 2bp step, and smoothed with a window size of 8 for the Hann function.

**Finding non-reference TE insertions from short reads**
Non-reference TE insertions were detected from Illumina data using TEBreak (https://github.com/adamewing/tebreak) with recommended parameters, apart from the following: -d tebreak/lib/hg38.chr.disctgt.txt -m tebreak/lib/hg38.chr.centromere_telomere.bed --min_sr_per_break 2 --skip_chroms chroms.txt. The file 'chroms.txt' was used to limit insertion calls to only canonical chromosomes (chr1-22, X, Y). TEBreak output was annotated with known non-reference insertions and filtered using the script provided (tebreak/scripts/general_filter.py). Insertion calls from TEBreak were compared against insertion calls from TLDR using the script "compare_tebreak_TLDR.py" included in the te-nanopore-tools repository. In this script, mappability of insertion sites (Supplementary Fig. 11) was determined via an index derived from mappability output from the GEM read mapper[15].

**Finding non-reference TE insertions from long reads**
We developed TLDR to analyse non-reference TE content from long reads. TLDR is available at https://github.com/adamewing/TLDR. While this study focuses on ONT long reads, TLDR can in principle use any accurately-aligned reads long enough to span TE insertions, including PacBio data. TLDR is a python application utilising multiprocessing that can complete analysis of a 30x long-read genome in under 1 hour (walltime) when utilising 32 cores. The operation of TLDR proceeds through two phases: clustering and insertion resolution. The required inputs consist of one or more sorted, indexed .bam files (recommended aligner is minimap2), a reference genome .fasta indexed with samtools faidx, and a set of reference TE sequences in fasta format (reference for human is included in distribution). Recommended options include --color_consensus for clear annotation of insertion features using ANSI colours and -p to specify a number of processes for multiprocessing. The results in this study were generated using parameters -e TLDR/ref/teref.human.fa -n TLDR/ref/nonref.collection.hg38.liftover.bed.gz --color_consensus -c chroms.txt -o all_extend --extend_consensus 20000 --detail_out. The file 'chroms.txt' was used to limit insertion calls to only canonical chromosomes (chr1-22, X, Y).

In the clustering phase (Supplementary Fig. 10a), clusters are seeded by identifying insertions (i.e. long indels) completely embedded in long reads. One read containing a completely embedded insertion bounded by a minimum and a maximum length (default 200-10000bp) is required to seed generation of a cluster. Additional reads are added to the cluster if they have an apparent breakpoint at either end of the seeding insertion, allowing for some ambiguity (default 200bp) around the junction. Nanopore reads can be arbitrarily long, bounded by the length of the chromosome, so one read can have membership in multiple clusters. In principle, this step could be further informed by input from more accurate short read mappings. As reads are allowed to be up to one chromosome in length, the clustering step can be parallelised on a per-chromosome basis (i.e. all chromosomes run at once).

The processing phase is parallelised on a per-cluster basis and begins with trimming clusters around the seeding insertion based on a user-specified flank size ($F$, default 500bp). For a given cluster each read is aligned against a set of reference TEs, required as input to TLDR, using an external program (exonerate with an affine:local model)[16] (Supplementary Fig. 10b). Up to three alignments at a minimum of 80% identity are reported for each read to allow for internal rearrangement of TEs versus the input reference (e.g. LINE-1 5′ inversions). Per-read alignments are assigned to groups which contain non-overlapping sets of alignments based on the best aligning reference TE. The group with the highest alignment score indicates the identity of the reference TE. Within each cluster $C$, only reads with alignments corresponding to this reference TE are used going forward. We refer to this subset as $C_{useable}$. At this point clusters meeting the minimum read count requirements are retained (default is 3 reads total with at least 1 read fully containing the insertion). Supporting reads are filtered by transforming the TE-aligning fraction of each supporting read into z scores and rejecting reads where abs(z) > 2. A cluster is rejected If a reference TE cannot be assigned or less than 50% of the reads in a cluster align to the majority reference TE.

A consensus sequence is generated for each cluster ($C_{useable}$) using multiple sequence alignment (MSA) via MAFFT[17] (Supplementary Fig. 10c). For each column in the MSA, the majority base (A,T,C,G, or gap) is returned where a gap majority can be overridden by two votes from one of (A,T,C,G) (i.e. two “A” bases would override a gap). Gaps are removed from the final consensus. This consensus sequence is then further refined through the following procedure. For each consensus sequence, reads from $C_{useable}$ are aligned back to the consensus sequence using minimap2 and the pileup (i.e. output from samtools mpileup -B -Q 1) is examined (Supplementary Fig. 10d). For each pileup column, the consensus sequence base is changed if greater than 50% of reads in the column vote for a different base with at least 3 votes for the same non-consensus base. For each column, if the pileup depth is less than 10% of the number of usable supporting reads in the cluster, the consensus base is not changed.

TE insertion breakpoints ($b_1 <= b_2$) are initially defined by fitting Gaussian mixture models (GMM) with 1 or 2 means (i.e. an insertion can have 1 or 2 breakpoints depending on the presence of TSDs or not) and choosing the best-fit model based on Akaike information criterion. These initial estimates are used to extract the insertion region from the reference chromosome from position $b_1$-$F$ to position $b_2$+$F$ (Supplementary Fig. 10e). The consensus is aligned against this reference sequence using minimap2 to refine breakpoint locations, and to annotate bases as aligned to the reference or part of the insertion (Supplementary Fig. 10f). All alignments (primary, and supplementary) are considered where the gap-compressed per-base divergence (i.e. “de” tag) is less than 0.12. Alignments are converted to a per-read profile where matched bases, inserted bases, and soft-clipped bases are encoded. These profiles are merged vertically to yield an overall mask where reference bases covered by a matched base are encoded as 1, and 0 otherwise. This mask is applied to the consensus, marking reference bases in upper case (A,T,C,G) and inserted bases in lower case (a,t,c,g). In each masked consensus, the longest inserted segment is aligned against the reference TEs using exonerate. The initial subfamily designation is allowed to change based on this new alignment. In cases where there is >1 unmapped segment, if the TE alignment spans both segments the segments are merged and intervening reference bases are presumed inserted. The refined breakpoints are used to define an initial TSD, which is then expanded to maximize the number of bases in the TSD. If the TSD is extendable, the TE alignment is repeated after TSD extension (Supplementary Fig. 10g).

If the user has requested detailed output (--detail_output), part of which is useful for assessing the methylation status of non-reference TE insertions (Supplementary Fig. 10h), additional per-insertion information is compiled in a directory named based on the input .bam file(s). This includes a per-insertion file containing information on supporting read mappings and a per-insertion file containing the consensus sequence. If the consensus sequence is extended (--extend_consensus), which is recommended for calling methylation on non-reference insertions, the consensus will be integrated into a larger segment of the reference genome sequence. The detailed output also includes a per-insertion, per-sample .bam file where the reads in the region defined by the extended consensus are aligned against the extended consensus, which includes the insertion sequence. Given the detailed output and a nanopolish indexed fastq (and associated fast5 files), non-reference methylation likelihoods can be obtained using nanopolish call-methylation[3] via the script included in scripts/TLDR_callmeth.sh. Additional auxiliary scripts are included for plotting and tabulating methylation data for non-reference insertions.

The output of TLDR is a tab-delimited file with columns as described in Supplementary Table 7. The consensus sequence output includes upper case characters representing reference bases and lower case characters representing the inserted sequence. The consensus can optionally be coloured (ANSI colours, viewable in compatible terminals via commands such as “cat” and “less -R”) via the --color_consensus command. If this is enabled, the inserted TE sequence will appear blue, inserted non-TE sequence (e.g. untemplated bases and transductions) will appear yellow and TSDs will appear red.

**Supplementary Table Legends**

**Supplementary Table 1:** Statistics for samples sequenced in this study. Note: sample representative identifiers were hc5413, CTRL-5413 normal hippocampus; he5413, CTRL-5413 normal heart; li5413, CTRL-5413 normal liver; hcc33nt, patient HCC33 non-tumour liver; hcc33t, patient HCC33 tumour.

**Supplementary Table 2:** Methylation statistics for reference TEs derived from ONT data. Tab **(a)** contains information for CTRL-5413 tissue samples and **(b)** contains information regarding the HCC33 liver tumour / non-tumour pair. Columns are as follows. Seg_chrom, seg_start, seg_end: position of the reference TE. Seg_name: subfamily name. Seg_strand: + or - indicating the orientation of the TE relative to the genome assembly. *_meth_calls: number of methylated CpG calls for each sample (i.e. nanopolish log likelihood ratio > 2.5). *_unmeth_calls: number of unmethylated CpG calls for each sample (i.e. nanopolish log likelihood ratio < -2.5). *_no_calls: number of ambiguous CpG calls for each sample (log likelihood ratio between -2.5 and 2.5). *_methfrac: fraction of non-ambiguous calls indicating methylation. Nearest_gene, gene_dist: nearest gene and distance between TE and said gene (0=intronic) based on Ensembl build 97 on hg38. Gene_type: derived from Ensembl annotation. Diff_sample (tab (a) only): indicates which sample is the most distant from the other two in terms of methylation fraction. Highest_meth (tab (b) only): indicates which sample has the highest methylation fraction. Fisher_p: Fisher's exact test p-value from comparison of methylation/demethylation counts between sample indicated in "diff_sample" (a) or "highest_meth" (b) and the other sample(s). Bonferroni_p: The value in "fisher_p" corrected for multiple testing. P-values are used here for ranking only and should not be taken as a measure of biologically relevant differences.

**Supplementary Table 3:** Output of TLDR run on all 5 ONT sequenced genomes. Tab **(a)** contains insertion calls passing all filters, **(b)** contains two known somatic L1Hs insertions specific to the HCC33 liver tumour sample[21], **(c)** contains insertion calls that do not pass one or more filters, **(d)** contains evidence for VNTR length variation derived from TLDR SVA-in-SVA calls. Column descriptions are as described in Supplementary Table 7.

**Supplementary Table 4:** Connections between insertions with transduced sequences and their likely individual (progenitor) element. Column descriptions are as follows. Chrom, Start, End, Family, Subfamily: see Supplementary Table 7. Ins_Source: indicates whether the apparent individual insertion is in the reference genome (REF) or corresponds to a known non-reference insertion (NONREF). Trans_loc: indicates whether the translocation is located on the 5′ or 3′ end of the insertion. individual: coordinates of the individual insertion in the reference genome (hg38). donor_family: annotation of individual elements. donor_source: "hg38" indicates the insertion was in the reference genome and annotated by repeatmasker, "uncatalogued" indicates no known reference or non-reference TE insertion is documented at the transduction mapping location. Other values indicate publications where non-reference insertions are catalogued. Notes: a free-form field for points of interest regarding the insertion or individual element.

**Supplementary Table 5:** Methylation statistics for selected reference TEs derived from ONT data. LINE-1 (L1), *Alu*, and SVA Insertions were considered if they met length thresholds and had at least 10 methylation calls. Tab **(a)** contains information for CTRL-5413 tissue samples and **(b)** contains information for the HCC33 liver tumour / non-tumour pair. Columns are as described for Supplementary Table 2 with the addition of the first column, seg_id, which is a UUID linking to Supplementary Table 3.

**Supplementary Table 6:** TEBreak output run on Illumina NovaSeq data from samples CTRL-5413 heart (he5413) and HCC33 non-tumour (hcc33nt). Tab **(a)** contains insertion calls passing all filters, tab **(b)** contains insertion calls that did not pass one or more filters. Columns are as described in the TEBreak documentation (https://github.com/adamewing/tebreak).

**Supplementary Table 7:** Tabs **(a)** and **(b)** provide column and filter descriptions, respectively, for TLDR output.
